# Demise of Marimermithida refines primary routes of transition to parasitism in roundworms

**DOI:** 10.1101/2022.02.15.480519

**Authors:** Alexei V. Tchesunov, Olga V. Nikolaeva, Leonid Yu. Rusin, Nadezda P. Sanamyan, Elena G. Panina, Dmitry M. Miljutin, Daria I. Gorelysheva, Anna N. Pegova, Maria R. Khromova, Maria V. Mardashova, Kirill V. Mikhailov, Vladimir V. Yushin, Nikolai B. Petrov, Vassily A. Lyubetsky, Mikhail A. Nikitin, Vladimir V. Aleoshin

## Abstract

Nematodes (roundworms) are ubiquitous animals commonly dominating in ecological communities and networks, with many parasites and pathogen vectors of great economic and medical significance. Nematode parasites are remarkably diverse in life strategies and adaptations at a great range of hosts and dimension scales, from whales to protozoan cells. Their life history is intricate and requires understanding to study the genomic, structural and ecological bases of successful transitions to parasitism. Based on analyses of rDNA for a representative sampling of host-associated and free-living groups, we dismiss the last higher-rank nematode taxon uniting solely parasitic forms (Marimermithida) and show that primarily marine parasitism emerged independently and repeatedly within only few free-living lineages. We re-evaluate the significance of some traditionally important phenotypic characters and report the phenomenon of dramatic adaptation to parasitism on very short evolutionary timescales. A cross-phylum character interpretation vindicates that non-intestinal (in-tissue or cavitary) host capture was likely a primary route of transition to truly exploitive parasitism (vs. intestinal commensalism) in roundworms, and extant nematode parasitoids (larval parasites) infesting the host body cavity or internal organs realise this primary lifestyle. Parasitism may have evolved in nematodes as part of innate pre-adaptations to crossing environmental boarders, and such transitions have been accomplished multiple times successfully in the phylum history.

## 1. Introduction

Nematodes (roundworms) are ubiquitous animals commonly dominating in ecological communities and networks. The majority are free-living inhabitants of marine sediments and soils, but manifold are parasites and pathogen vectors with often a global economic and medical impact, which colonised a great range of hosts at all dimension scales, from whales to protozoan cells. Parasitism in nematodes is a richly diverse phenomenon from the perspectives of biology, ecology and the evolutionary routes of transition to parasitic lifestyle. Understanding life history of successful transitions from free-living ancestors is necessary to study the genomic, structural and environmental bases of parasitism in general biology, medical science and pest control. In the currently embraced phylogenomic framework^1–5^, there have been speculated a series of independent transitions to plant (at least 3), non-vertebrate (10) and vertebrate (5) parasitism of various form and extent across the entire phylum^6–8^. Some other views develop a more diversified scenario of at least 20 independent transitions to insect parasitism alone^9^. Among the three stem nematode lineages, the richest repertoire of parasitic lifestyles is observed among the Chromadoria clade, followed in ranking by Dorylaimia towards only a few known cases in Enoplia. The overwhelming majority of parasitic nematodes belong within lineages of terrestrial or limnic ancestry^6,10^, while the descendent marine parasites are accordingly implied secondary invaders to the sea. Current knowledge derives roundworms from a marine ancestor inhabiting sea sediments of the Ediacaran—Cambrian at about 600— 500 Ma (e.g., ^refs.11,12^), which also gains support from solid phylogenomic evidence (e.g., ^ref.13^) of their kinship with well-documented mid-Cambrian marine tardigrades (e.g., ^refs.14,15^). Among all nematodes, the Enoplia stem assertively gains basal position in phylogenetic analyses of protein-coding (a representative sampling in ^refs.2,4,5^), SSU rDNA data (large-scale phylum-inclusive assays, e.g. ^ref.10^) and in a comparative framework of early embryogenesis^11,16,17^, nervous system architecture^11^, gonad and sperm cell morphology and ultrastructure^18–20^. Enoplians comprise diverse mainly marine free-living forms abundant in sediments towards the deep sea, where they also contribute to the core of nematode communities^21,22^. However, the only commonly recognised enoplian parasites belong to the Triplonchida clade, which is terrestrial, brackish or freshwater-inhabiting. This seemingly definite mapping of parasitic associations on the nematode phylogeny stipulates their exclusive on-land ancestry and highlights the importance to identify and explore cases of primarily marine parasitism in roundworms to enable hypotheses on the routes, bases and predispositions of parasitism in this most speciose and abundant group of metazoan life.

Prevailing and most impactive nematode parasites in the sea belong to the chromadorian Spirurina clade inferred as clade III in ^ref.1^, where they infest a wide diversity of mostly vertebrate hosts (bony fish, sharks and marine mammals). Marine spirurians supposedly derived from on-land parasitic ancestors via invertebrate vector transmission to definitive hosts in a freshwater-anadromous-marine sequence (e.g.,^refs.11,23^) between the Devonian—Carboniferous and early Mesozoic (origin of modern-type spirurians)^12,23^. A major lifecycle trait they share with other nematode parasites is the free-to-host phase transition at juvenile/larva stage 3 (J3, third of four pre-adult stages in nematode lifecycle), which may also carry a phoretic or diapausing function^24^. Most chromadorian parasites invade, and dorylaimian parasitoids (early-cycle larval parasites) exit the host as J3. Meanwhile, a major distinguishing feature between spirurian and non-spirurian marine parasites is definitive host habitat. The latter comprise sparse instances across Enoplia, Dorylaimia and Chromadoria and never invade the gut but exclusively the body cavity and its structures in a wide range of mostly benthic non-vertebrate hosts (sea urchins, starfishes, annelids, molluscs, priapulids, other nematodes or even testate gigantic protozoans^25–29^). Alike on-land forms, marine parasitoids dwell as juveniles in the body cavity or internal organs exhausting the host individual to death and abandon the corpse into the milieu to become free non-feeding reproductive adults. Alike spirurians, enoplian and dorylaimian parasites may possess a gigantic size (up to tens of cm vs. about 1–20 mm in their free-living relatives) and strong fecundity associated with adaptations in the female reproductive system^27,28^. These characters are traditionally assigned evolutionary significance, albeit without a clear appraisal in a molecular phylogenetic context. I this study, we integrated original sequences (i.a. on rare primarily marine parasites) with various-source rDNA data (i.a. environmental metagenomic and amplicon surveys) available to combine host-associated and free-living nematode groups into a representative phylogeny. It was used to interpret evidence of life traits, morphology and ecology in a non-formal approach (due to a limited statistical power of formal reconstruction methods for a few complex characters). In the discussion, we attempt to explain the combination of traits in more basal primarily marine vs. more derived limnic-terrestrial lineages in order to suggest (i) the ancestral mode of host capture, lifecycle and habitat type, (ii) an evolutionary scenario that produced the diversity of modern parasitic lifestyles, (iii) to assess evolutionary significance of important parasitism-related traits and (iv) identify putative pre-adaptations to successful parasitism in baseline nematode organisation and ecology.

Historically, the thin grade of marine non-spirurian parasites were classified in morphology-based systems in the orders Mermithida (an established on-land taxon of Dorylaimia), Marimermithida, Benthimermithida and Rhaptothyreida (all *incertae sedis*^11,30– 32^). Despite rarity, these groups spread widely across the world ocean, from the tidal zone down the depth of 5.2 km^ref.30^. They are hard to collect and captured only occasionally, often upon off-field examination of formalin-fixed hosts. The first molecular data on selected individuals initiated re-shifting and fitting-in the enigmatic aberrant taxa into a mature phylogenetic context. Thus, the spectacular giant parasite of sea urchins *Echinomermella matsi*, initially subsumed as mermithid, was relocated to another stem lineage, in the crown of Enoplida^33^. Another aberrant marine parasite *Nematimermis enoplivora*, a minute-scale dweller of body cavity of other nematodes^34^, is verified as member of Dorylaimia for the first time with molecular data in present study. The affinity of rare deep-sea trophosome-feeding benthimermithids to the chromadorian stem in larval morphology^35^ was confirmed in molecular phylogenetic trees^36–38^. Upon discovery in deep-see corers, bizarre standing-out Rhaptothyreida were considered alternatively in kinship with Mermithida^39^, Marimermithida or Benthimermithida^40^ based on the trophosome character derived from vestigial mouth and alimentary tract. To date, rhaptothyreids have never been found inside a host but have a juvenile morphology strikingly different from putative adult, i.a. in minute cephalic sensilla, the absence of feather-like amphids and presence of a stylet-like structure in buccal cavity, which may suggest their larval parasitism, with free-living adults and in-host juveniles^41^ (noting that latest stages observed in this group so far may still be immature individuals; A.V.T, unpublished). The phylogenetic placement of rhaptothyreids is confined recently within the order Enoplida in rDNA trees, although with a fuzzier certainty therein^42^. The last and major traditional group of marine parasitic nematodes is Marimermithida. The genus *Marimermis* was erected by Rubtzov and Platonova^25^ basing on three novel species extracted from body cavity of starfish or found freely in inter-to upper subtidal sediments. The authors have surmised marimermithids to be larval parasites (parasitoids) of marine invertebrates, displaying an unusual combination of typically parasitic (e.g., up to 15 cm-long body, trophosome-transformed gut) and marine free-living traits (e.g., cephalic and abundant body sensilla). United later with the superficially similar benthimermithid *Trophomera* and another three genera, *Marimermis* established the aberrant taxon Marimermithida^43^, which until 2021 remained in essence the last stronghold of higher-rank nematode systematics for its uncertainty in phylogenetic placement among roundworms. An affinity of marimermithids to free-living marine Enoplida has been suggested^11^ but with no confident appraisal due to the inaccessibility of biological samples for molecular analyses. First molecular data on the marimermithid genus *Aborjinia* recently revealed its membership within the Enoplia stem, although without a reappraisal of the whole taxon monophyly^44,45^. Here, we report molecular evidence on two representatives of Marimermithida, as well as on other rarely captured marine nematode parasites, and discuss the nematode life strategies and evolutionary adaptations to parasitism within the framework of resulting phylogeny.

## 2. Material and methods

### 2.1. Material collection and sampling sites

Specimens of *Marimermis maritima* were discovered in coelomic cavity of sea urchin *Strongylocentrotus polyacanthus* (Figure 1 A, 1). The hosts were diver-collected in the northwest Pacific Ocean near the coast of island Matua (48.1° N, 153.2° E) in August 2016. The *Aborjinia* sp. specimen was found in a priapulid *Priapulus caudatus* (Figure 1 B, 1). The host was collected from 5,349–5,352 m depth in the northwest Pacific Ocean, Kurile-Kamchatka Trench, with an Agassyz trawl during the KuramBio deep-sea expedition (R/V Sonne, SO-223) in August 2012. *Nematimermis enoplivora* was discovered inside the nematode *Enoplus communis* (Figure 1 C, 1). Hosts were collected with muddy inter-to subtidal sediment near the White Sea Biological Station of Moscow State University (Kandalaksha Bay, White Sea; geographic locale is same hereafter) during summer 2009. Individuals of phanodermatid K2 and *Camacolaimus* spp. were found in-test of the foraminifer *Reophax curtus* collected by subtidal trawling during summers 2010–2015. Specimens isolation, processing and free-living nematodes collection are detailed in electronic supplementary material.

**Figure 1.**
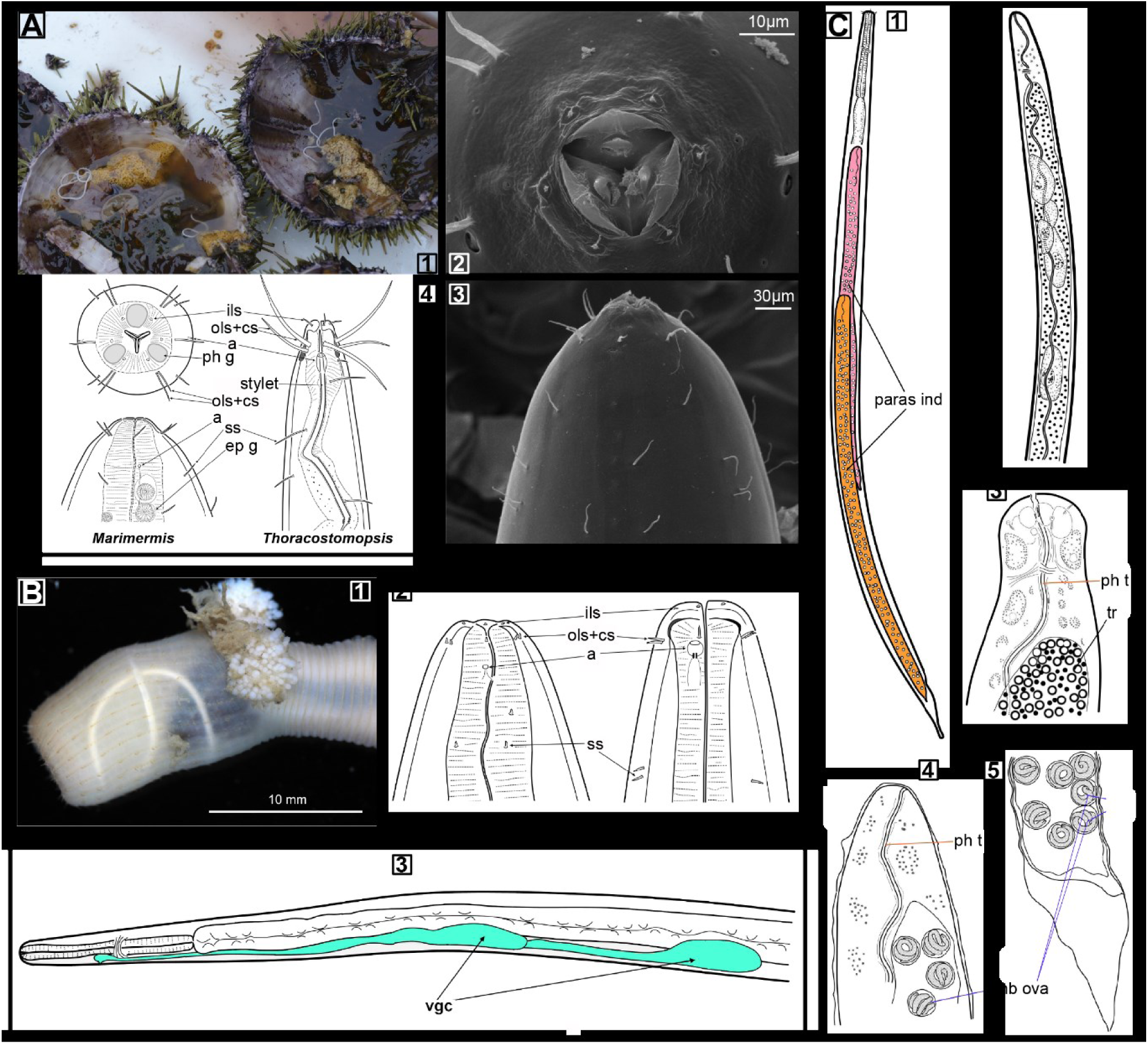
Cavitary marine nematodes **A** – *Marimermis maritima*. 1 – Live specimens exposed in open host, sea urchin *Strongylocentrorus polyacanthus*. 2, 3 – Cephalic end apically and laterally, respectively, scanning electron microphotograph; pharyngeal gland outlets on lips, abundant irregular somatic setae. 4 – Generalised anterior ends juxtaposed in *M. maritima* and *Thoracostomopsis barbata* (Enoplida, Thoracostomopsidae). **B** – *Aborjinia* sp. 1 – Juvenile occupying introvert of *Priapulus caudatus* (courtesy by Dr. A.S. Maiorova, NSCMB FEB RAS). 2 – Generalised anterior ends juxtaposed in *Aborjinia* sp. and *Leptosomatum* sp. (Enoplida, Leptosomatidae). 3 – Anterior body with two-celled ventral secretory-excretory gland, a distinctive genus feature. **C** – *Nematimermis enoplivora*. 1 – Two mature parasitoid individuals within host body of nematode *Enoplus communis* (Enoplida, Enoplidae). 2, 3 – Anterior body and head of immature parasitoid retrieved from body cavity of *E. communis*. 4, 5 – Anterior and tail regions of mature parasitoid from *E. communis*; trophosome resorbed, embryos with larval stylet inside egg shells, nested retained exuvia (old molted cuticles). Abbreviations: a – amphid, emb ova – embryonated eggs, ep g – epidermal glands, ils – inner labial sensilla, ols+cs – joint crown of outer labial and cephalic setae, paras ind – parasitoid individuals, ph g – pharyngeal glands, ph t – pharyngeal tube, ss – somatic setae, stylet – spear-shaped armature, tr – trophosome, vgc – ventral gland cells.

### 2.2. Dataset construction and phylogenetic analysis

DNA/RNA extraction and sequencing, data processing, phylogenetic pipeline, sequence origins and accession IDs are detailed in electronic supplementary material. Original data were obtained by Sanger or NGS sequencing (depending on the object; NCBI BioProject PRJNA772260, NCBI sequence IDs provided on Figure 2) and incorporated in a study-specific various-origin (including metagenomic and amplicon eDNA surveys) dataset accommodating a representative cross-phylum diversity of host-associated and free-living groups. The rDNA gene alignments were constructed using RNA secondary structure-aware algorithms, concatenated and used for maximum-likelihood (ML) and Bayesian (BI) tree inference. Phylogenetic hypotheses were verified in appropriate ML-based statistical tests of topologies. [The partitioned alignment is available at submission as electronic supplementary file.]

**Figure 2.**
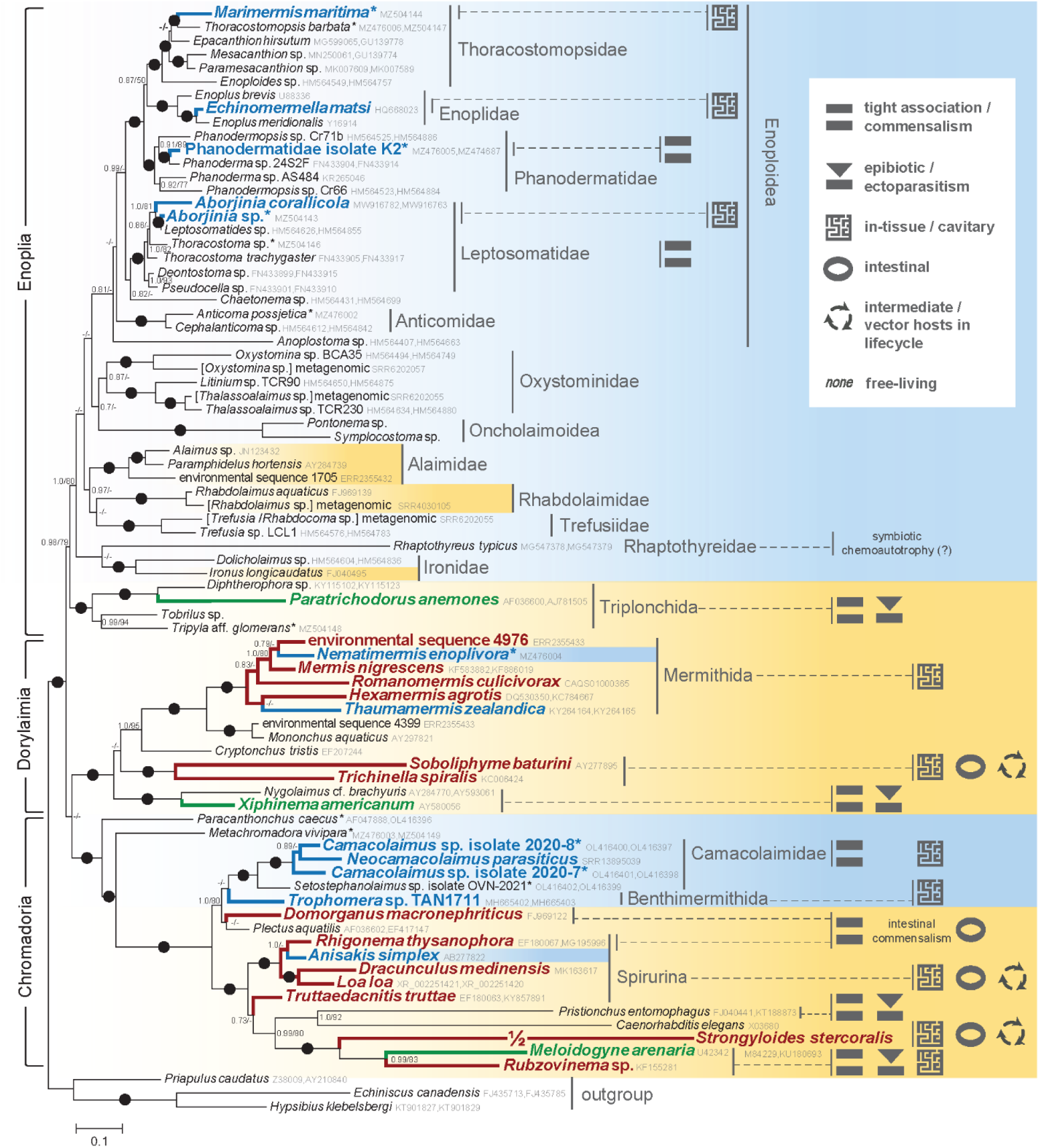
Bayesian tree of Nematoda based on combined SSU, 5.8S and LSU rDNA data spanning the three major clades (Enoplia, Dorylaimia and Chromadoria), with extended sampling of marine Enoplia and a selection of representative host-associations (**in bold**) for the clades. Parasitism-relevant life traits are mapped as pictograms, explained in legend and addressed in Discussion. Traits apply either to particular sampled species or more broadly to a subclade it represents (according to general knowledge of typical large subclades). Pictogram series denote traits co-occurrence either within subclade or individual lifecycle. Habitat is colour-coded to generally typify subclades (e.g., as per ^ref.10^), blue – marine, sandy – brackish/limnic/terrestrial. Marine animal parasites/associates are in dark blue, brackish/limnic/terrestrial – in terracotta, plant parasites/associates – in green. Nodes mismatched in BI and ML topologies are unlabelled. Otherwise, labels contain BI posterior probabilities (left) and ML bootstrap support (right). Values <0.7/70 replaced with dashes (–). Bipartitions with support 1/100 dotted (●). BI posterior probabilities are calculated across GTR+*Г* parameter space in 3 M generations, ML bootstrap support estimated under GTR+F+G16 model in 100 replicates. NCBI accessions appended to taxon names, original sequences and assemblies marked with asterisk (*). Scale bar: substitutions per site.

## 3. Results

### 3.1. Specimen identification

Individuals of *Marimermis maritima* (Figure 1, A), *Aborjinia* sp. (Figure 1, B) and *Nematimermis enoplivora* (Figure 1, C) were expert-identified in this study to species or genus level based on morphometry and descriptive data in light microscopy. The phanodermatid K2 specimen was selected from among few juveniles extracted from foraminiferan hosts captured in a single sample and identified with certainty to family level as Phanodermatidae. Detailed morphology data and verification are available for expert evaluation in electronic supplementary material. All other nematode specimens used for DNA sequencing originally in the study were expert-identified to species or genus level live prior to fixation.

### 3.2. Phylogenetic analysis

ML and BI multi-gene rDNA-based reconstructions produced congruent topologies in most areas except for selected deeper nodes of the nematode tree (Figure 2, supplementary Figure S1; result reproducible with SSU and LSU partitions separately, Figures S2–S3). Because the ML method of finding a single best tree is more susceptible to model misspecification and heterogeneity-associated biases, we depict the BI tree on Figure 2 and leave the mismatched nodes unlabelled for node support. The two topologies are not discriminated in ML-based statistical hypothesis testing (electronic supplementary material), overall well supported and conform to the established phylum phylogeny^3–5^. The host-associated species of the study are separated in different nodes with high support, except for *Rhaptothyreus typicus*. The latter branch is very long and may be consistently misplaced across phylogeny due to associated inference artefacts, which has already been reported in^42^. Of key relevance to the discussion is the absence of its validated monophyly with any of the host-associated lineages in statistical tests. Other study species of Enoplia were unambiguously placed in different traditional taxonomic families, although united within single superfamily (Enoploidea) of the order Enoplida. The traditional order Marimermithida is paraphyletic: *Marimermis maritima* groups within Thoracostomopsidae, and *Aborjinia* sp. – within Leptosomatidae. The partial SSU and LSU genes of *Leptosomatides* sp. from ^ref.3^ are nearly identical with those of *Aborjinia* sp. from this study; they were obtained for a free (off-host) individual captured in marine sediment near the coast of California at a 2,694 m depth, which therefore likely represents a misidentified specimen of *Aborjinia* that had abandoned the host. *Echinomermella matsi* falls inside the genus *Enoplus* as member of the family Enoplidae, the closest clade to Thoracostomopsidae. The foraminifer-associated phanodermatid K2 groups within Phanodermatidae in sistership with *Phanoderma* sp. Outside Enoplia, the same-habitat *Camacolaimus* spp. specimens form a confident clade with the cavitary annelid parasite *Neocamacolaimus parasiticus* within the monophyletic assemblage of Camacolaimidae (a family of Chromadoria), which also includes the deep-sea cavitary invertebrate parasite *Trophomera* sp. *Nematimermis enoplivora* is confidently inferred in the crown of the order Mermithida within Dorylaimia, thus corroborating an earlier morphology-based view^34^ and revealing its secondary intrusion to the sea and primary on-land ancestry.

## 4. Discussion

Our phylogenetic analyses of rDNA cistron genes and a rigor hypothesis testing unequivocally resolve the relationships of several non-spirurian marine host-associated taxa (Figure 2) and clearly demonstrate multiple origins of primarily marine parasitism in roundworms. The benthimermithid *Trophomera* finds its place within Chromadoria as a deviant parasitic marine representative of the mainly freshwater-terrestrial plectid lineage, in accord with ^36–38^, in sistership with a peculiar assemblage of minute-size nematodes (Camacolaimidae) associated with foraminiferan protozoans or parasitising coelom of microscopic annelids. With Enoplia, it was rather unexpected that all spots of emergent parasitism within the stem are confined to single traditional superfamily Enoploidea otherwise consisting entirely of free-living forms. The position of Rhaptothyreida is established outside Enoploidea within the order Enoplida, in agreement with the original study^42^. *Rhaptothyreus typicus* appears very long-branched in rDNA trees and requires a focused systematic analysis, with exploring more taxon sampling and parameter variation until additional molecular markers are available. An ML-based hypothesis testing reveals its undefined placing with no confident affinity to mutually non-monophyletic host-associated lineages (electronic supplementary material). Despite within-lifecycle morphological disparity suggesting larval parasitism in this enigmatic worm, in-host findings of rhaptothyreids are presently unknown. Their trophosome cells are stuffed with putative bacterial particles^46^, which altogether may suggest a non-parasitic but aberrant chemoautotrophic lifestyle, similar to vestimentiferan and pogonophoran annelids and alike in the mouthless nematodes *Astomonema* and *Parastomonema*^32^.

The morphology-based affinity of *Nematimermis enoplivora* (Figure 1 C) is confirmed within the Dorylaimia stem in a traditional order uniting on-land invertebrate parasitoids^34^. It assertively suggests its recent trespass to the sea from an on-land ancestor parasitising a terrestrial or brackish-water host. There are two morphological descriptions that may explicate this trajectory in mermithids by the far-flung cases of infecting a supra-to intertidal arthropod (*Thaumamermis zealandica* of sandhopper^47^, with molecular evidence in ^ref.48^; ref. to Figure 2) towards inhabiting an unknown host in deep sea (*Thalassomermis megamphis*^49^). We provide a molecular-verified report of truly marine parasitism in Mermithida, which may be considered an extant event of host-borne intrusion to the sea, thus supporting the hypothetic scenario of vector-borne transition to marine vertebrate hosts realised early in the chromadorian evolution by ancestral spirurians^11,23^.

An unusual instance of primarily marine parasitic-like nematode associations reported in this study is phanodermatid K2. It represents an unidentified species of the family Phanodermatidae in our molecular analyses and the first report of cavitary association in this otherwise free-living enoplian group. Phanodermatid K2 thus complements a cohort of peculiar associates of the large (about 1 mm size) unicellular testate agglutinated foraminifer *Reophax curtus* predominating in meiobenthic communities of the White Sea^50,51^. *Reophax curtus*, among other benthic foraminifers, is known to provide a niche for miniature-scale parasitism in the chromadorian family Camacolaimidae^52–54^ that invade about 10% of the foraminifer population. The worms dwell within a vacuole or interstitially in a cavity between cytoplasm and the test and probably can leave or reinvade the host at variant stages via the ostium or other wall pores. A non-mutualistic nature of this association is evident from a relatively low invasion rate, cytoplasm lesion and a protective host response by enclosing the intruder within a proteinaceous capsule. Alike in camacolaimids, some phanodermatid specimens in our samples exhibited peculiar exaggerated pharyngeal glands, perhaps due to enhanced secretion in-test of the foraminifer (observations by D.I.G., unpublished, ref. to electronic supplementary material). Nutrients are likely absorbed in both solid (alimentarily) and solute (transcuticularly) form, as entails from a residual poor gut content in microscopy, also suggesting an exploitive nature of this alliance. Among the invasive camacolaimid species discovered, only few are obligately associated with the foraminifer and not found in sediments (A.V.T., unpublished), which reveals a transient nature of this cavitary association between the free-living state towards facultative and permanent host capture. An independent instance of same association in a marine enoplian illustrates the capacity of cavitary parasitism as a universal transition route that is being exploited in evolution multiple times successfully at a minute-dimension scale as well. The realisation of this scenario even at a short evolutionary timescale gains support in our present finding of a confident assemblage uniting protist-associated forms (*Camacolaimus* spp. isolates) and a true coelomic parasite of microscopic benthic polychaetes (*Neocamacolaimus parasiticus*^55^).

The position of the giant sea urchin parasite *Echinomermella matsi* within the otherwise free-living genus *Enoplus*^33^ is re-confirmed. This species has no direct relationships with mermithids or other known marine parasitic nematodes. An in-genus sistership of radically diverged lifestyles and phenotypes is rather unprecedented and represents a natural model system to study genomic predispositions to parasitism and generate insights into particular mechanisms underlying host capture and in-host survival. This type of knowledge is key to understanding evolutionary transition pathways and prerequisite to develop drug and vaccine candidates against known or potential disease agents. Commonly, free-to-host transitions occur over long timescales and accumulate genomic disparity, which obscures comparisons between the parasitic and free-living model counterparts. Recent research in this field has benefited from two narrow systems of emerging or facultative parasitism in three strongyloidid genera (Chromadoria clade IV)^56^ and the genus *Caenorhabditis* (Chromadoria clade V)^57^. Two different evolutionary scenarios have been exposed, an evident genomic reduction in the strong commensal *Caenorhabditis bovis* (at the expense of G-coupled receptor-coding genes involved in sensory systems and cell-to-cell communication) and a negligible loss but massive gene families expansion (mostly in astacin-like and SCP/TAPS immunomodulatory proteins), as well as their invasion-specific upregulation, in facultative and obligate strongyloidid parasites. A close relatedness to a free-living genome enabled strong inference of selective gene family duplications in both models and transcriptomic/proteomic evidence of their direct association with in-host survival towards a proposal of operonic control of parasitism-associated genes in *Strongyloides* species. Despite rarity of the *E. matsi* parasite, the *Echinomermella–Enoplus* complex may represent a promising model to direct future research in this field owing to its unique phylogenetic compactness combined with biological and structural disparity that far surpass the mentioned systems. *Echinomermella matsi* is a cavitary parasite and obviously represents a separate, primarily marine route of emergent parasitism to complement in evidence the two different scenarios in strongly commensalic primarily free-living soil rhabditids and intestinal strongyloidid parasites adopting a more intricate and probably descended vector-lost strategy of percutaneous J3 larva invasion and in-gut migration inside the host. A free-living component of the model is already available with genomic and transcriptomic data on *Enoplus brevis*^58^. Meanwhile, this instance of outbreaking parasitism entailing radical change in phenotype and lifestyle within a narrow lineage in not unique among nematodes. As we exemplify below with our findings, similar spots of explosive adaptation are discovered within the Enoploidea superfamily and may in fact be more common across the phylum. This observation suggests that such an extent of life trait plasticity during adaptive evolution is inherent in roundworms as a phenomenon endemic among other Metazoa.

A solid statement of this study is paraphyly of the traditional order Marimermithida and its confident redistribution on nematode phylogeny. This inference implies a higher-rank taxonomic restructuring of a part of the nematode system and dismisses the last major taxonomic division that united marine parasites alone. Our analyses reject the hypotheses of marimermithid affinity with the order Mermithida or anywhere close within the Dorylaimia stem (as assumed, e.g., in ^ref.40^ and later in ^ref.59^) but distribute them between the two distinct traditional families within the Enoploidea superfamily in Enoplia. Accordingly, the peculiar life traits uniting the former taxon and shared with dorylaimian mermithids (larval cavitary parasitism in invertebrates, enlarged size and intense fecundity) thus need be reinterpreted as potentially convergent. Although the enoplian origin of marimermithids has been grounded in pharyngeal and sensory morphology^25,35,43^ (ref. to Figures 1 A, 2–4, B, 2), the inference of *Marimermis maritima* within the ramified crown of Enoplia in family Thoracostomopsidae is unexpected and was not anticipated before. Thoracostomopsids are marine free-living nematodes known as predators and scavengers^60^. They retain normal nematode anatomy with some particular structures in buccal armoury evidently used to grip food particles, e.g., a smaller nematode prey. None of thoracostomopsid species was known as parasitic or associated with any host. One particular character in free-living thoracostomopsids is, however, remarkable to functionally resemble that in a cohort of non-related parasitic nematodes. *Thoracostomopsis barbata*, the closest free-living relative of *M. maritima* in current dataset, represents the sole thoracostomopsid genus possessing a long eversible spear in the mouth cavity (Figure 1 A, 4). The spear is a joint structure compounding elements along entire buccal cavity^61^ and used apparently for piercing food objects, albeit the genus diet is unknown. This spear structure is unique among free-living nematodes^62^. Spear-like structures or stylets of various origin engaged in penetrating the host are known from the invasive larvae of on-land dorylaimian vertebrate parasites Dioctophymatida and Trichinellida, invertebrate parasitoids Mermithida, on-land chromadorian insect- and plant-parasitising Tylenchina and certainly from marine chromadorian invertebrate-infesting Benthimermithida^29^ as part of the associations-rich camacolaimid assemblage, to also mention nematomorphs (hairworms) outside Nematoda^63^. These structures are not homologous across taxa. Invasive larvae in *Marimermis* remain undiscovered, but we posit them to possess a spear-like structure resembling that in *Thoracostomopsis*. Another noticeable character of Thoracostomopsidae are the pronounced sophisticated epidermal glands in some genera. The glands in *T. barbata* have a fine fringe radial striation^62^ resembling the elaborate rosette-like epidermal organs in *Marimermis*^25,29^. Although their functional load is unstudied, active levels of epidermal secretion implied by their presence are likely inherent in transcuticular transmission and permeability processes critical in parasitic in-host survival and feeding and/or prerequisite to crossing environmental boarders. Cuticular and secretion lability is considered in some views among the baseline characters to enable the spectacular potential of nematodes to conquer new niches^10^.

*Aborjinia* sp. is inferred separately from *M. maritima* within Enoploidea as an internal branch of the family Leptosomatidae, in line with ^44,45^. It also shares with its relatives some elaborate glandular structures in tail region that suggest enhanced secretion^28,30^ typical in marine free-living nematodes, as well as a peculiar two-celled ventral secretory-excretory gland or renette^28^ (Figure 1 B, 3). This organ is mono-celled in the majority of free-living and parasitic forms, with a rare two-celled deviation described in two *Leptosomatum* species^64^. In contrast to *M. maritima* and its relatives, leptosomatid nematodes possess more bland uniform morphology that is less transformed in *Aborjinia*: minute nearly papilloid sensory sensilla, normal muscular pharynx, no elaborate armature in mouth cavity and normal cellular midgut allowing food ingestion and alimentary feeding. They are non-predacious large-bodied (up to 20 mm and more) forms considered giants among marine free-living nematodes. With this complex of features, Leptosomatidae is the sole enoplian family reported to have strong associations with cnidarian anthozoans^65^ to the extent of true in-tissue parasitism of *A. corallicola* recently discovered inside stolon body of a cold-water octocoral and placed on molecular trees^44^. The association with coral is clearly epibiotic in emergence and exemplifies a pertinent case of gradually transforming lifestyle to capture host. Some leptosomatids are known to facultatively dwell within sponges and even feed from these hosts^66^, thus revealing prerequisites to adopting a parasitic lifestyle. In-sponge environment is close to marine but its compartmentalisation may predispose physiological and behavioural adaptations to facultative towards permanent cavitary parasitism.

The confident allocation of primarily marine parasitic and parasitic-like associations on nematode phylogeny demonstrates their independent emergence and allows comparison of structural and life traits in these vs. free-living and more derived groups. It can be observed that permanent host capture can be attained via transient associations of various form and extent realised in tight communities close to epibiotic towards occupancy of internal spaces in sourcing for nutrients and protection. Such communities likely emerge and maintain temporally over entire lifecycle of the animal, predisposing infestation early in the cycle at a larval stage. Direct colonisation of gut proceeded naturally in this scenario via adaptations to strongly commensalic connections with host. Conversely, truly exploitive associations of surviving on host’s own resources emerged in an alternative evolutionary pathway. We surmise that cavitary larval host capture in direct lifecycle was likely a primary route of transition to true parasitism in roundworms, and extant parasitoids (larval parasites) infesting the host body cavity or internal organs realise the primary parasitic lifestyle. More complex strategies involving intermediate or vector hosts and definitive intestinal stages can be derived via natural extension of within-lifetime capture beyond one host along contemporary environmental food chains towards new resources and more effective dispersal. Such transmissions have evolved in advanced nematode parasites of the trichinellid–dioctophymatid lineage in Dorylaimia and historic Secernentea group in Chromadoria, especially in spirurians and strongyloidids (Figure 2). Nowadays they commonly exhibit peculiar in-host migration patterns of the infective larva to reach the gut destination^67^, which may adequately be explained by secondary fallout of the primary cavitary stage in lifecycle. The hypothetic cavitary scenario gains direct support from occasional exquisite Early Devonian nematode fossils preserved in stomatal chambers of an early land plant^68^. The finding demonstrates family clusters of worms, including eggs, juveniles and adults, in substomatal and cortical cavities, as well as interstitially inside plant tissue, with indications of on-site reproduction and perhaps a facultative semi-bacterivorous nature of association. The fossil quality allowed its putative assignment close to an extant marine subtaxon of Enoplia due to a barely changed free-living morphology. It is a grounded speculation that the primary nematode body plan and physiology already provided the prerequisites for successful colonisation of miscellaneous inner environments, and such transitions have been realised multiple times successfully in the phylum history. Transient associations can be facilitated at minor or no morphological adaptation, as exemplified above with strongly commensalic soil rhabditids or by the case of even stronger intestinal commensalism of debris-inhabiting chromadorian *Domorganus*^69,70^, with some of its species foraging in-gut of oligochaetes in same manner its closest relatives do in soil. We do observe that a stylet-like armoury in mouth cavity is often associated with evolutionarily advanced host capture conducing to true parasitism in non-related groups of both animal and plant-parasitic nematodes. This structural feature distinguishes lineages enriched with parasitic or parasitic-like lifestyles and having a non-gut (in-tissue or cavitary) nature of such associations, even at a closer evolutionary scale. For instance, the diverse camacolaimid assemblage in Chromadoria unites stylet-bearing forms and includes both minute transient protozoan in-test associates and larger true animal coelomic parasites (including deep-sea benthimermithids), whilst the stylet-less sister *Domorganus* realises a different strategy of strong intestinal commensalism (Figure 2). Unlike this morphological precondition of true parasitism, oversize, strong fecundity and non-alimentary feeding (truly transcuticular or trophosome-based) typical of parasitic forms can be considered secondary adaptations to permanent host capture likely gaining advantage in nutrient-rich but temporary habitats requiring dispersal in lifecycle. It can be envisaged that roundworms are probably unique metazoans that originated as a higher-rank lineage already pre-equipped with the necessary toolkit in morphology and physiology to enable effective expansion across environmental niches, including parasitism. We should expect and pursue particular achievements from focusing on the genomic and physiological bases of nematode parasitism to explain its higher incidence within some natural lineages uniting mostly free-living forms (e.g., Enoploidea in Enoplia reported in this study or Camacolaimidae in Chromadoria), as well as clarify preconditions to selectively occupying non-intestinal environment at the earliest primary stage of transmission. This line of research also holds perspective to explore limits of animal phenotypic adaptation by capitalising on peculiar natural model systems of compact parasitic—free-living species complexes that we begin to discover among the diversity of roundworms.

## Acknowledgements

The authors are thankful to the administration and staff of White Sea Biological Station (Moscow State University) for providing excellent cooperative working environment, research facilities and infrastructure, Tatyana S. Miroliubova (Severtsov Institute of Ecology and Evolution) for assistance in laboratory works, Elena R. Alekhina and Lyudmila N. Ryzhonkina (Belozersky Institute of Physicochemical Biology, Moscow State University) for involvement in field sampling, Prof. Dr. Andrey V. Adrianov and Dr. Anastassya S. Maiorova (A.V. Zhirmunsky National Scientific Center of Marine Biology, FEB RAS) for the priapulid host identification and image, the organisers and participants of the 20th and 21st Russian Geographical Society’s Kamchatka-Kurile expeditions to island Matua (middle Kurile Islands) in 2016 and 2017 for providing sampling resources and to the team of Podvodremservis business enterprise (Petropavlovsk-Kamchatsky) for managing diving equipment.

## Electronic supplementary material

**1. Supplementary results and methods**

