## Supplementary Material for "Demise of Marimermithida refines primary routes of transition to parasitism in roundworms"

<u>**Electronic supplementary material**</u>

**1. Supplementary figures**

**2. Molecular data origin and availability**

**3. Supplementary Materials and Methods**

3.1. Material extraction, processing and free-living nematodes collection

3.2. DNA/RNA extraction and sequencing

3.3. Dataset and phylogenetic pipeline

3.4. Statistical ML tests of phylogenetic hypotheses

**4. Biological material identification**

4.1. Specimen identification in *Marimermis maritima* (Figure 1, A)

4.2. Specimen identification in *Aborjinia sp.* (Figure 1, B)

4.3. Specimen dentification in phanodermatid K2 isolate

4.3.1. Female morphology of Phanodermatidae gen. sp.

4.3.2. Microdrawing of female Phanodermatidae gen. sp.

**5. Supplementary references**

### 1. Supplementary figures

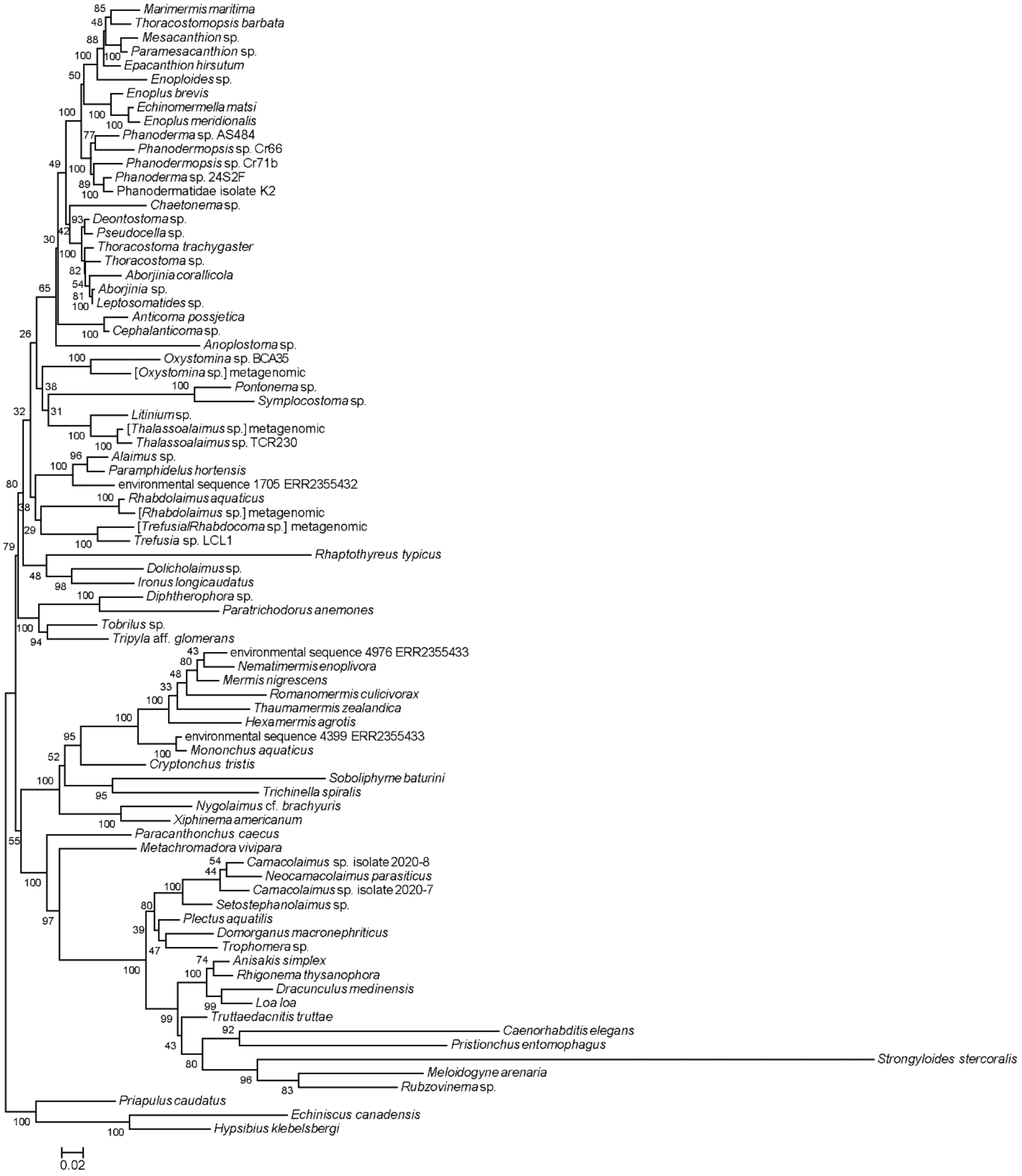


**Figure S1**. ML tree of Nematoda based on combined SSU, 5.8S and LSU rDNA data. Nodes labelled with bootstrap support values estimated under GTR+F+G16 model in 100 replicates. Scale bar: substitutions per site.

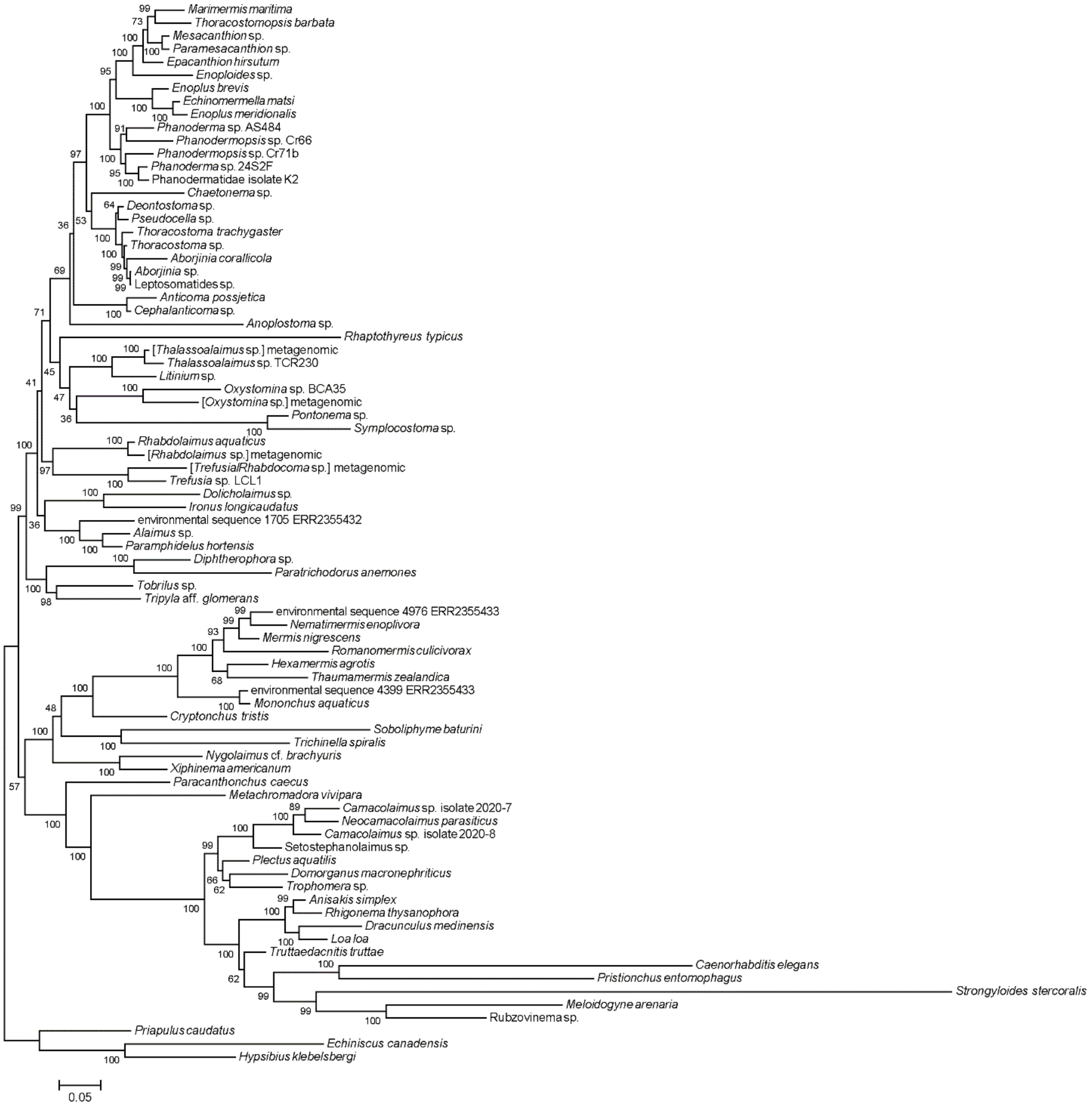


**Figure S2**. Bayesian tree of Nematoda based on SSU rDNA data. Nodes labelled with posterior probabilities (%) calculated across GTR+*Γ* parameter space in 3 M generations. Consensus topology obtained with 50% burn-in. Scale bar: substitutions per site.

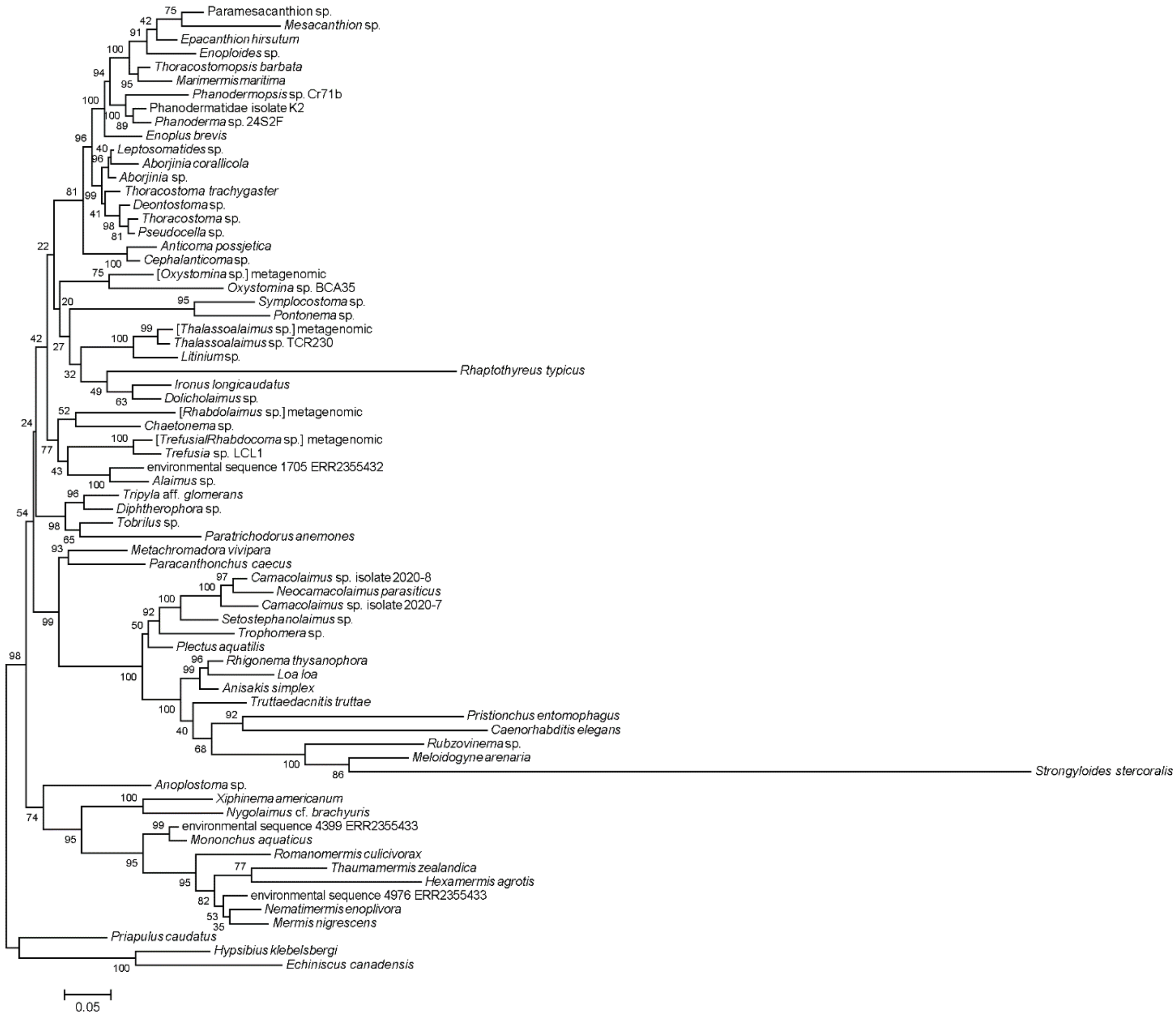


**Figure S3**. Bayesian tree of Nematoda based on LSU rDNA data. Nodes labelled with posterior probabilities (%) calculated across GTR+$\Gamma$ parameter space in 3 M generations. Consensus topology obtained with 50% burn-in. Scale bar: substitutions per site.

#### 2. Molecular data origin and availability

Sequences obtained originally in the study. NCBI GenBank accession IDs provided for genes. Raw NGS data available in NCBI BioProject PRJNA772260

<table>
<tr><th>Taxonomic unit</th><th>Method</th><th>BioSample</th><th>SSU</th><th>5.8S</th><th>LSU</th></tr>
<tr><td>Aborjinia sp.<br>OVP-2021 isolate KKT</td><td>Sanger</td><td></td><td colspan="3">MZ504143</td></tr>
<tr><td>Anticoma possjetica</td><td>Sanger</td><td></td><td>MZ476002</td><td></td><td></td></tr>
<tr><td>Camacolaimus sp.<br>OVN-2021 isolate 2020-7</td><td>Sanger</td><td></td><td>OL416401</td><td></td><td>OL416398</td></tr>
<tr><td>Camacolaimus sp.<br>OVN-2021 isolate 2020-8</td><td>Sanger</td><td></td><td>OL416400</td><td></td><td>OL416397</td></tr>
<tr><td>Marimermis maritima</td><td>Sanger</td><td></td><td colspan="3">MZ504144</td></tr>
<tr><td>Metachromadora vivipara</td><td>NGS</td><td>SAMN22371214</td><td>MZ476003</td><td colspan="2">MZ504149</td></tr>
<tr><td>Nematimermis enoplivora</td><td>Sanger</td><td></td><td>MZ476004</td><td></td><td></td></tr>
<tr><td>Paracanthonchus caecus<br>isolate Pc-OVN-2021</td><td>Sanger</td><td></td><td></td><td></td><td>OL416396</td></tr>
<tr><td>phanodermatidae gen. sp.<br>OVP-2021 isolate K2</td><td>Sanger</td><td></td><td>MZ476005</td><td></td><td>MZ474687</td></tr>
<tr><td>Setostephanolaimus sp.<br>isolate OVN-2021</td><td>Sanger</td><td></td><td>OL416402</td><td></td><td>OL416399</td></tr>
<tr><td>Thoracostoma sp.<br>OVP-2021 isolate C-Tsp</td><td>NGS</td><td>SAMN22371215</td><td colspan="3">MZ504146</td></tr>
<tr><td>Thoracostomopsis barbata</td><td>NGS</td><td>SAMN22371216</td><td>MZ476006</td><td colspan="2">MZ504147</td></tr>
<tr><td>Tripyla aff. glomerans<br>OVP-2021 isolate C-Tg</td><td>NGS</td><td>SAMN22371213</td><td colspan="3">MZ504148</td></tr>
</table>

Sequences originally assembled from third-party NGS data

| Taxonomic unit | BioProject | SRA |
|---|---|---|
| *Neocamacolaimus parasiticus* | PRJNA707491 | SRR13895039 |
| [*Oxystomina* sp.] metagenomic | PRJNA415343 | SRR6202057 |
| [*Thalassoalaimus* sp.] metagenomic | PRJNA415343 | SRR6202055 |
| [*Trefusia / Rhabdocoma* sp.] metagenomic | PRJNA415343 | SRR6202055 |
| [*Rhabdolaimus* sp.] metagenomic | PRJNA336658 | SRR4030105 |

Third-party NCBI GenBank gene accession IDs

<table>
<tr><th>Taxonomic unit</th><th>SSU</th><th>5.8S</th><th>LSU</th></tr>
<tr><td>Aborjinia corallicola voucher S11306-2</td><td>MW916782</td><td></td><td>MW916763</td></tr>
<tr><td>Alaimus sp. SK-2012</td><td></td><td></td><td>JN123432</td></tr>
<tr><td>Anisakis simplex</td><td colspan="3">AB277822</td></tr>
<tr><td>Anoplostoma sp. BUS21</td><td>HM564407</td><td></td><td>HM564663</td></tr>
<tr><td>Caenorhabditis elegans</td><td colspan="3">X03680</td></tr>
<tr><td>Cephalanticoma sp. TCR141</td><td>HM564612</td><td></td><td>HM564842</td></tr>
<tr><td>Chaetonema sp. NAR6</td><td>HM564431</td><td></td><td>HM564699</td></tr>
</table>

| | | | |
|---|---|---|---|
| *Cryptonchus tristis* | EF207244 | | |
| *Deontostoma* sp. 1S2F8 | FN433899 | | FN433915 |
| *Diphtherophora* sp. Shahrood | KY115102 | | KY115123 |
| *Dolicholaimus* sp. TCR114 | HM564604 | | HM564836 |
| *Domorganus macronephriticus* strain DoGaMac2 | FJ969122 | | |
| *Dracunculus medinensis* isolate PDB18-022 | MK163617 | | |
| *Echiniscus canadensis* isolate Tar14 | FJ435713 | | FJ435785 |
| *Echinomermella matsi* | HQ668023 | | |
| *Enoploides* sp. DBA1 | HM564549 | | HM564757 |
| *Enoplus brevis* | U88336 | | |
| *Enoplus meridionalis* | Y16914 | | |
| *Epacanthion hirsutum* | MG599065 | | GU139778 |
| environmental sequence 1705 | ERX2404076 / ERR2355432 | | |
| environmental sequence 4399 | ERX2404077 / ERR2355433 | | |
| environmental sequence 4976 | ERX2404077 / ERR2355433 | | |
| *Hexamermis agrotis* | DQ530350 | | KC784667 |
| *Hypsibius klebelsbergi* voucher HD005 | KT901827 | | KT901829 |
| *Ironus longicaudatus* | FJ040495 | | |
| *Leptosomatides* sp. TCR192 | HM564626 | | HM564855 |
| *Litinium* sp. TCR90 | HM564650 | | HM564875 |
| *Loa loa* | XR_002251421 | | XR_002251420 |
| *Meloidogyne arenaria* | U42342 | | |
| *Mermis nigrescens* isolate Mjuv | KF583882 | | KF886019 |
| *Mesacanthion* sp. 2 TJP-2019 Nem.77 | MN250061 | | |
| *Mononchus aquaticus* | AY297821 | | |
| *Nygolaimus* cf. *brachyuris* JH-2004 | AY284770 | | AY593061 |
| *Oxystomina* sp. BCA35 | HM564494 | | HM564749 |
| *Paracanthonchus caecus* | AF047888 | | |
| *Paramesacanthion* sp. 2 AAS-2018 isolate SB101 | MK007609 | | MK007589 |
| *Paramphidelus hortensis* | AY284739 | | |
| *Paratrichodorus anemones* | AF036600 | | AJ781505 |
| *Phanoderma* sp. 24S2F | FN433904 | | FN433914 |
| *Phanoderma* sp. AS484 | KR265046 | | |
| *Phanodermopsis* sp. Cr66 | HM564523 | | HM564884 |
| *Phanodermopsis* sp. Cr71b | HM564525 | | HM564886 |
| *Plectus aquatilis* | AF036602 | | EF417147 |
| *Pontonema* sp. | Smythe et al. 2019 | | |
| *Priapulus caudatus* | Z38009 | AY210840 | |
| *Pristionchus entomophagus* | FJ040441 | | KT188873 |
| *Pseudocella* sp. 3S26E8 | FN433901 | | FN433910 |
| *Rhabdolaimus aquaticus* | FJ969139 | | |
| *Rhaptothyreus typicus* | MG547378 | | MG547379 |
| *Romanomermis culicivorax* | CAQS01000365 | | |
| *Rubzovinema* sp. EIK-2013 | KF155281 | | |
| *Soboliphyme baturini* | AY277895 | | |
| *Strongyloides stercoralis* | M84229 | | KU180693 |

| | | | |
|---|---|---|---|
| *Symplocostoma* sp. | Smythe et al. 2019 | | |
| *Thalassoalaimus* sp. TCR230 | HM564634 | | HM564880 |
| *Thaumamermis zealandica* | KY264164 | | KY264165 |
| *Thoracostoma trachygaster* | FN433905 | FN433917 | |
| *Tobrilus* sp. | Smythe et al. 2019 | | |
| *Trefusia* sp. LCL1 | HM564576 | | HM564783 |
| *Trichinella spiralis* | KC006424 | | |
| *Trophomera* sp. TAN1711 | MH665402 | | MH665403 |
| *Truttaedacnitis truttae* | EF180063 | | |
| *Xiphinema americanum* | AY580056 | | |

Original assemblies by Smythe et al. 2019[ref.1] are available at https://figshare.com/s/4c8e501714dbd5be1be8

#### 3. Supplementary Materials and Methods

*3.1. Material extraction, processing and free-living nematodes collection*

Host individuals of sea urchin *Strongylocentrotus polyacanthus* were opened unfixed, and specimens of *Marimermis maritima* collected from coelomic cavity and preserved with 96% ethanol for DNA extraction.

The host individual of *Priapulus caudatus* was dissected unfixed, and *Aborjinia* sp. specimen retrieved from introvert haemocoel onboard of the vessel. The nematode was divided in three parts; anterior and posterior ends were fixed with 10% buffered seawater formalin for light microscopy. The middle portion was subdivided in two; one part was fixed with 100% ethanol and stored at –20°C, another part – with DESS[2] and stored at room temperature; both parts were used for DNA extraction.

Host specimens of *Enoplus communis* were extracted from muddy inter- to subtidal sediment by repeated decantation and fine-sieving. The *Nematimermis enoplivora* parasite was retrieved live from body cavity of scalpel-punctured hosts, sliced and fixed with 96% ethanol for DNA extraction.

Foraminifers *Reophax curtus* were cavitation-extracted from muddy subtidal sediment of trawl catch and fixed with DESS. Individuals of Phanodermatidae and *Camacolaimus* spp. were isolated manually from fixed tests under stereo microscopes.

Specimens of *Paracanthonchus caecus*, *Thoracostomopsis barbata*, *Pseudocella trichodes*, *Thoracostoma* sp. and *Metachromadora vivipara* were sampled from subtidal sandy sediments near the White Sea Biological Station of Moscow State University, *Tripyla* sp. – from freshwater detrital debris of the Chernaya river bed in August 2018 (Kandalaksha Bay, White Sea). Nematodes were extracted by decantation-sieving and fixed with RNAlater stabilisation solution (Thermo Fisher Scientific, USA) for total RNA extraction. Individuals of *Anticoma possjetica* were obtained from bivalve *Crenomytilus grayanus* clusters collected at a 10 m depth near the Vostok Biological Station of A.V. Zhirmunsky National Scientific Centre of Marine Biology in August 2004 (Vostok Bay, Sea of Japan); worms fixed with 96% ethanol for DNA extraction.

*3.2. DNA/RNA extraction and sequencing*

Individual nematodes were lysed in 20 µl proteinase K-containing buffer[3]. Total DNA was extracted with NucleoSpin Tissue XS kits (Macherey-Nagel, Germany) following a manufacturer protocol. PCR amplification of partial SSU (18S) rDNA, complete ITS regions and partial LSU (28S) rDNA was conducted with specific primers[4,5] using Encyclo PCR chemistry (Evrogen, Russia). PCR cycling: primary denaturation at 95°C for 5 min, cycle denaturation at 95°C for 30 s (40 cycles), annealing at 55°C for 30 s, extension at 72°C for 3 min, final extension at 72°C for 5 min. PCR products were agarose gel-purified with Cleanup Mini kits (Evrogen, Russia). Amplicons were sequenced directly with an Applied Biosystems 3730 DNA Analyzer (Thermo Fisher Scientific, USA). Original transcriptomic data was generated in *Metachromadora vivipara*, *Pseudocella trichodes*, *Thoracostomopsis barbata*, *Thoracostoma* sp. and *Tripyla glomerans*. Total RNA from nematode samples was extracted using Arcturus PicoPure RNA Isolation Kits (Thermo Fisher Scientific, USA). Transcriptomic libraries for *P. trichodes*, *T. barbata* and *Thoracostoma* sp. were prepared with the TruSeq protocol (Illumina, USA). In *M. vivipara* and *T. glomerans*, cDNA was synthesised with SMART-Seq v4 Ultra Low Input RNA Kits (Takara Bio, France). Libraries were sequenced on an Illumina HiSeq 4000 system, generating 16–37 M 150-bp paired-end reads for each sample. Original raw NGS data are deposited as NCBI BioProject PRJNA772260, rDNA cistron assemblies and original Sanger sequences are deposited in NCBI GenBank.

*3.3. Dataset and phylogenetic pipeline*

Illumina reads were processed with Trimmomatic[6] and *cutadapt*[7] tools to remove adapter sequences and assembled using SPAdes[8] with *k*-mer values 77 and 127. Contigs corresponding to rDNA operons were extracted from SPAdes assemblies using BLAST[9]. Fragmented rDNA operons were merged by overlapping contigs or contig termini, and the resulting operon assemblies were examined for errors by read mapping with Bowtie2[ref.10] and the mapping inspection in Tablet[11]. Additional rDNA sequences were obtained from the GenBank *nr*, *wgs*, SRA and *figshare* repositories (see above). Individual alignments of SSU, 5.8S and LSU rDNA were prepared with MAFFT[12] using Q-INS-i RNA secondary structure-aware algorithm. Variable indel-rich hairpin regions were additionally realigned by X-INS-i algorithm incorporating pairwise structural alignment by MXSCARNA[13] and using conserved flanking hairpins as alignment anchors. For tree inference, the alignments were masked with trimAl[14] using the built-in automated trimming heuristic (-automated1). The alignment length was 1703 bp in SSU, 152 bp in 5.8S and 2390 bp in LSU rDNA. Three gene alignments were manually concatenated (90 taxa total) and used for tree inference as partitioned supermatrix or separately. Bayesian phylogenetic inference (BI) was performed using MrBayes 3.2.6[ref.15] with two runs (nst=6 ngammacat=16 rates=invgamma), three partitions (SSU, 5.8S, LSU), 3,000,000 generations and 50% burn-in. Average standard deviation of split frequencies was 0.4 on run completion. Runs converged across all parameters with PSRF[16] estimate 1.0. Maximum likelihood (ML) trees with bootstrap support were estimated in 100 replicates (–b 100) using IQ-TREE 1.6[17]. Best model was selected with ModelFinder (–m MF[18]) on topology fixed as BI consensus (–t). The best model initially selected according to BIC and AIC across partitions was GTR+F+G4; it was refined further with ModelFinder (–mset GTR+F) by explicitly testing rate heterogeneity types (–mrate) as G4, G6, G8, G10, G12, G14, G16 and G18. The GTR+F+G16 model was ultimately selected and used in downstream analyses. Phylogenetic trees were visualised with MEGA 6.0[ref.19]. Alternative hypothesis testing was performed according to the approximately unbiased (AU[ref.20]) and expected likelihood weight (ELW[21]) tests implemented in IQ-TREE 1.6. Hypotheses were formalised as topological constraints and used in ML inference with IQ-TREE under the model parameters pre-estimated as described above and 10,000 RELL replicates (–zb 10000) for bootstrap support approximation[22]. The constraint hypotheses and their related confidence values in a test tree set containing consensus ML and BI trees are detailed below in Section 3.4.

*3.4. Statistical ML tests of phylogenetic hypotheses*

ML-based hypothesis testing for monophyletic placement of the nematode lineages under study was performed as follows. The lineages with parasitic or other host-associated lifestyle were tested in pairwise combinations for exclusive monophyly against other nematode taxa (as test hypotheses) in a tree set containing all test hypotheses, including BI and ML consensus trees. Given the overall high-supported consensus trees, we reduced test combinatorics outside the primary subjects in Enoplia to only verity *Nematimermis enoplivora* for exclusion from Mermithida and Dorylaimia (hypotheses `N_OUT_MER` and `N_OUT_DOR`, respectively). Test hypotheses were formalised as topological constraints (ref. to table below) and used in ML inference with IQ-TREE 1.6. Approximately unbiased (AU, $p < 0.05$) and expected likelihood weight (`c-ELW`) statistical tests were used as implemented in IQ-TREE with 10,000 RELL replicates (`-zb 10000`) and `GTR+F+G16` model pre-selected. Non-rejected hypotheses and respective confidence values are **in bold**. Abbreviations are self-explaining and provided for clarity; Nematoda, Dorylaimia, Mermithida = the rest of dataset relative to the explicitly constrained taxa in each hypothesis.

All tests did not discriminate between the ML and BI trees. All pairwise hypotheses were rejected except for the grouping of *Rhaptothyreus typicus* with *Echinomermella matsi*. The latter exclusive monophyly is not rejected at a lower-boundary *p*-value in AU, rejected in ELW test and discords with the ML/BI topologies, which assertively suggests an uncertain volatile positioning of *R. typicus* and requires a separate methodological effort and/or further molecular evidence. The affinity of *N. enoplivora* to the outer tree with respect to both Mermithida and Dorylaimia is rejected, thus confirming its placement as a crown mermithid.

| Test hypothesis | AU *p*-value | c-ELW |
|---|---|---|
| **Bayesian** consensus | **0.518 +** | **0.498 +** |
| **Maximum likelihood** consensus | **0.625 +** | **0.476 +** |
| AAM ((Aborjinia corallicola,*Aborjinia* sp.,*Marimermis maritima*),Nematoda); | $6.26\times10^{-45}$ – | $6.35\times10^{-113}$ – |
| AAP ((Aborjinia corallicola,*Aborjinia* sp.,phanodermatid K2),Nematoda); | $5.42\times10^{-46}$ – | $1.76\times10^{-104}$ – |
| AAE ((Aborjinia corallicola,*Aborjinia* sp.,*Echinomermella matsi*),Nematoda); | $6.69\times10^{-6}$ – | $8.02\times10^{-55}$ – |
| AAR ((Aborjinia corallicola,*Aborjinia* sp.,*Rhaptothyreus typicus*),Nematoda); | 0.00802 – | 0.00379 – |
| MP ((*Marimermis maritima*,phanodermatid K2),Nematoda); | $2.14\times10^{-71}$ – | $1.67\times10^{-70}$ – |
| ME ((*Marimermis maritima,Echinomermella matsi*),Nematoda); | $6.09\times10^{-5}$ – | $2.51\times10^{-39}$ – |
| MR ((*Marimermis maritima,Rhaptothyreus typicus*),Nematoda); | 0.0166 – | 0.00447 – |
| PE ((phanodermatid K2,*Echinomermella matsi*),Nematoda); | $7.81\times10^{-116}$ – | $1.92\times10^{-22}$ – |
| PR ((phanodermatid K2,*Rhaptothyreus typicus*),Nematoda); | $1.89\times10^{-5}$ – | $9.41\times10^{-8}$ – |
| **ER ((*Echinomermella matsi,Rhaptothyreus typicus*),Nematoda);** | **0.0861 +** | $1.5\times10^{-8}$ – |
| AAMR ((Aborjinia corallicola,*Aborjinia* sp.,*Marimermis maritima,Rhaptothyreus typicus*),Nematoda); | $1.27\times10^{-5}$ – | $3.11\times10^{-114}$ – |
| N_OUT_DOR ((Dorylaimia), *Nematimermis enoplivora*, Nematoda); | $9.27\times10^{-5}$ – | $5.62\times10^{-165}$ – |
| N_OUT_MER ((Mermithida), *Nematimermis enoplivora*, Nematoda); | 0.0142 – | 0.0178 – |

#### 4. Biological material identification

Detailed morphology and verification data for specimen identification in *Marimermis maritima*, *Aborjinia* sp. and phanodermatid K2.

*4.1. Specimen identification in* Marimermis maritima *(Figure 1, A)*

The specimen was identified to species level based on morphology and descriptive data from [23,24].

*4.2. Specimen identification in* Aborjinia *sp. (Figure 1, B)*

The specimen of *Aborjinia* sp. was identified to genus level based on morphological and metric data. Main measurements: L = ca. 30 mm; a = 60; b = 20; c = 74; c′ = 1.4. Body diameters at level of: second head sensilla circle = 109 µm; amphid = 126 µm; nerve ring = 250 µm; cardia = 290 µm; anus = 290 µm. Maximum diameter = ca. 500 µm. Body whitish, opaque, cylindrical, slightly narrowed towards anterior end. Tail short, in shape of rounded cone. Internal organs visible hardly due to fleshy body walls consisting of very developed hypodermal chords and thick somatic musculature. Lateral hypodermal chords very wide; nuclei of numerous chordal cells visible. Total number of hypodermal chords not determined. In anterior end, muscular envelope having (in addition to longitudinal) diagonal and transversal muscle fibres. Somatic sensilla sparsely distributed, short, cylindrical, ca. 2.5 µm length, associated with hypodermal chords. Metanemes not found. Nerve ring situated at ca. 650 µm from head end. Caudal gland opening visible at terminal tail tip. Cephalic sensilla arranged in two circles (6 + 10). First circle represents six labial disc-shaped papilla (extremely minute) located at lip bases. Second circle consists of short cylindrical setae, four pairs of lateromedian (of them, first ca. 3 and second ca. 4–5 µm length) and two lateral setae ca. 3 µm length. Amphidial aperture small, pore-like (ca. 1–2 µm diameter), with clear prominent rim, situated at ca. 50 µm from head end. Canalis amphidialis hardly visible. Mouth opening surrounded by three lips (two subventral and one dorsal). Pharynx muscular, cylindrical. Three pharyngeal gland ducts visible in anterior pharynx, two subventral opening at terminal pharynx (ca. 5 µm from head apex) and dorsal – at ca. 10 µm posterior head apex. Cardia not seen. Intestine with visible internal lumen. Rectum hardly visible, lacking thick cuticular walls. Anus in shape of transversal slit. Ventral excretory gland opening visible on ventral side at ca. 520 µm posterior head end. Excretory gland comprises long duct and two large cellular bodies. Anterior- and posteriormost cellular bodes laying ca. 3.3 and 4.7 mm from head end. Vulva and gonads not found.

The examined juvenile definitely satisfies the morphological descriptive criteria of the family Marimermithidae. Among the distinctive features are: larval parasitic lifestyle, overall habitus, head sensillar pattern and location, amphid structure, three mouth lips, presence of three well-developed pharyngeal glands opening very close to mouth edge[24–27]. Very short or papilloid cephalic sensilla and a two-celled excretory gland with a long duct opening anterior to nerve ring are all characteristic of the genus *Aborjinia* Özdikmen 2010 (pro *Australonema* Tchesunov et Spiridonov 1985, junior homonym of *Australonema* Tassell 1980)[25–27]. Accordingly, the examined juvenile should be attributed to the genus *Aborjinia*.

*4.3. Specimen dentification in phanodermatid K2 isolate*

Specimen K2 was a juvenile selected for sequencing from a collection containing several juvenile individuals and one mature female retrieved from the foraminifer *Reophax curtus*. The specimens had slim fusiform bodies, smooth cuticle and discernible dark oculi typical of the free-living enoplid family Phanodermatidae. Some specimens exhibited peculiar exaggerated pharyngeal glands, perhaps due to enhanced secretion during in-test association with foraminifers, and a residual gut content suggesting a non-symbiotic exploitive nature of the association. The female and juveniles could not be identified with certainty to species or genus level and are hereby designated Phanodermatidae gen. sp. (fam. Phanodermatidae Filipjev, 1927; order Enoplida Filipjev 1929).

*4.3.1. Female morphology of Phanodermatidae gen. sp.*

Body fusiform, with smooth cuticle. Length 3,043 µm, middle diameter 66 µm. Cephalic setae in two closely spaced circles (4 + 6), 10 and 7 µm length. Amphids small, slit-shaped, ca. 4 µm. Body diameter at amphids 15 µm. Amphideal fovea prolongation not definable. Oral cavity minute to indefinable. Three protrusions visualised in stoma. Renette small-celled, with pore opening in anterior body end, duct continuing along pharynx. Pharynx muscular with prominent transverse striation, anterior third containing nerve ring with large accompanying assemblage of neuronal bodies. Posterior pharynx contains large glandular extension with three large pharyngeal gland cell bodies, two latero-ventral and one dorsal. Body diameter at this section 73 µm. Pharyngeal glands markedly exaggerated, open anteriorly into pharynx. Gut reveals distinct cell boundaries and central lumen discernible throughout and containing nutritive content. Anus ventral posterior. Body diameter at anal orifice 56 µm. Ovaries homodromous. Vulvar opening approximately middle-body. Caudal gland cell bodies posterior, with homogeneous cytoplasm, open at tail tip.

*4.3.2. Microdrawing of female Phanodermatidae gen. sp.*

Left – general view, scale bar 300 µm. Right – cephalic end, scale bar 20 µm.

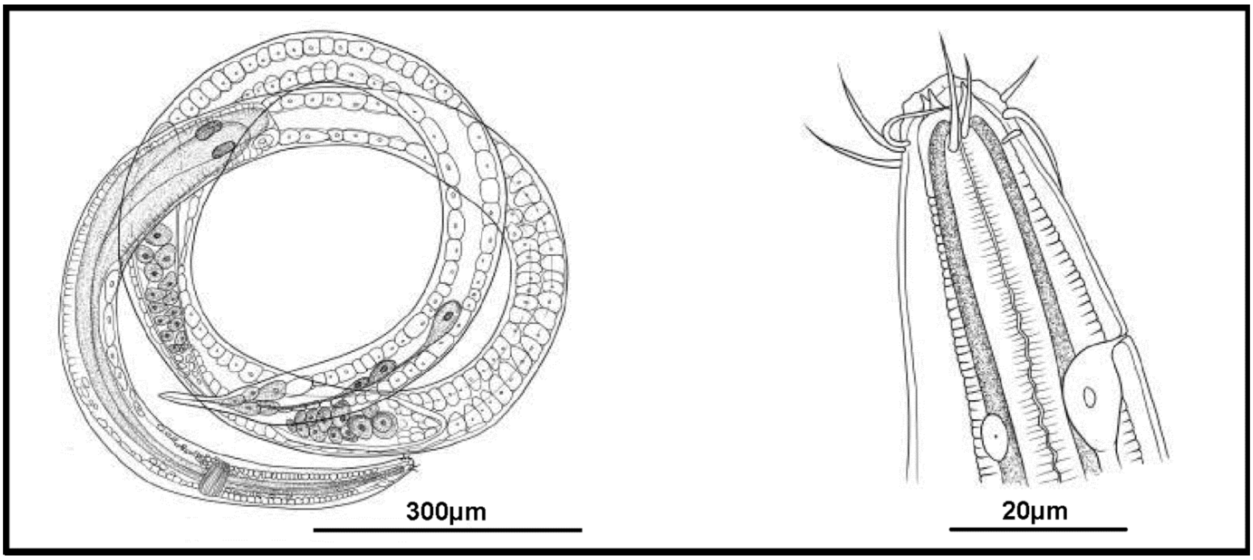
